# DLO Hi-C Tool for Digestion-Ligation-Only Hi-C Chromosome Conformation Capture Data Analysis

**DOI:** 10.1101/764332

**Authors:** Ping Hong, Hao Jiang, Weize Xu, Da Lin, Qian Xu, Gang Cao, Guoliang Li

**Author notes:** Correspondence to G.L. or G.C. These authors contributed equally to this work (Ping Hong, Hao Jiang, Weize Xu, Da Lin).

## Abstract

**Background:** It is becoming increasingly important to understand the mechanism of regulatory elements on target genes in long-range genomic distance. 3C (Chromosome Conformation Capture) and its derived methods are now widely applied to investigate genome organizations and gene regulation. Digestion-Ligation-Only Hi-C (DLO Hi-C) is a new technology with high efficiency and effective cost for whole-genome chromosome conformation capture.

**Results:** Here, we introduce DLO Hi-C Tool, a flexible and versatile pipeline for processing DLO Hi-C data from raw sequencing reads to normalized contact maps and providing quality controls for different steps. It includes more efficient iterative mapping and linker filtering. We applied DLO Hi-C Tool to different DLO Hi-C datasets, and demonstrated its ability of processing large data in multi-threading.

**Conclusions:** DLO Hi-C Tool is suitable for processing DLO Hi-C and in situ DLO Hi-C datasets. It is convenient and efficient for DLO Hi-C data processing.

## Background

Recently, there is increasing evidence that higher-order chromatin structures play an important role in gene expression and regulation [1]. In order to dissect the three-dimensional (3D) structure of the chromatin, chromosome conformation capture (3C) method [2] depending on DNA proximity ligation [3] was proposed in 2002. With the rapid progression of high-throughput sequencing technology and bioinformatics, various derivative methods were developed to study the physical interactions for different needs, such as Hi-C [4], ChIA-PET (Chromatin Interaction Analysis by Paired-End Tag Sequencing) [5], DNase Hi-C [6], in situ Hi-C [7] and Capture Hi-C [8]. Hi-C is a powerful method to investigate the genome-wide all-to-all long-range chromatin interactions and topologically associating domains (TADs) [9], and has contributed to great achievements in understanding the principle of biological functions from the perspective of 3D chromosomal structure.

Although Hi-C has greatly advanced our understanding of the 3D organization of genomes, it is limited with high randomly ligated DNA noise, high cost and complex experimental procedures. To overcome the limitations of Hi-C method, we developed a simple, cost-effective and low-noise method, Digestion-Ligation-Only Hi-C (DLO Hi-C) [10]. The key advantages of DLO Hi-C are as follows. Firstly, the main steps of DLO Hi-C method are simplified with only two rounds of digestion and ligation. Secondly, it is convenient to estimate the signal-to-noise ratio from different combinations of linkers, which is not available for other Hi-C methods. Thirdly, the linker-ligated fragments are purified by cost-effective polyacrylamide gel electrophoresis (PAGE) instead of biotin labeling and pulling down steps, which reduces the cost in the experiment. Fourthly, simultaneous digestion and ligation steps can effectively reduce the percentage of re-ligation reads. Lastly, because of the delicate design of specific nucleotide barcodes, double digestion is convenient to be used for an early quality-control step to examine the ratios of ligation compositions before sequencing to evaluate the quality of DLO Hi-C library. By applying two rounds of digestion and ligation, approximately 80-bp DLO Hi-C DNA fragments were generated which contains a 40-bp complete linker and a pair of 20-bp interacting genomic DNA fragments. These DLO Hi-C fragments were subjected to high-throughout sequencing. The pair of the mapped genomic DNA fragments will uncover chromatin interactions. Self-ligation products from the two ends of the individual DNA fragments or re-ligation products (the paired-end DNA interaction information derived from two adjacent DNA fragments) just take a small proportion in the uniquely mapped reads. As far as we know, there is no pipeline appropriate for DLO Hi-C data. In order to facilitate the analysis of DLO Hi-C data, here we present an easy-to-use and versatile pipeline called ‘DLO Hi-C Tool’ to process DLO Hi-C data from raw sequencing reads to normalized contact maps.

In this report, we provide a detailed description of the design and implementation of DLO Hi-C Tool. By applying DLO Hi-C Tool to different DLO Hi-C datasets, we have demonstrated its efficiency and effectiveness. Quality control at different steps of the DLO Hi-C analysis is supported in our pipeline and will be displayed in the final report files.

## Design and Implementation

DLO Hi-C Tool pipeline (Fig. 1) begins with sequencing reads from DLO Hi-C library. There are four main steps: pre-processing of raw sequencing reads, reads aligning and filtering, noise reduction and paired-end reads classification, and interaction visualization. It is available for users to modify any modules with flexibility. Information can be transported through file system between steps, so that users can resume the process if the program is interrupted by accident. In this program, most of functions was code by Java and compiled into jar package. There is no need for users to install the program, just to ensure that the version of Java is no less than 1.8 and the dependent software (BWA and python) have been installed correctly.

**Fig. 1.**
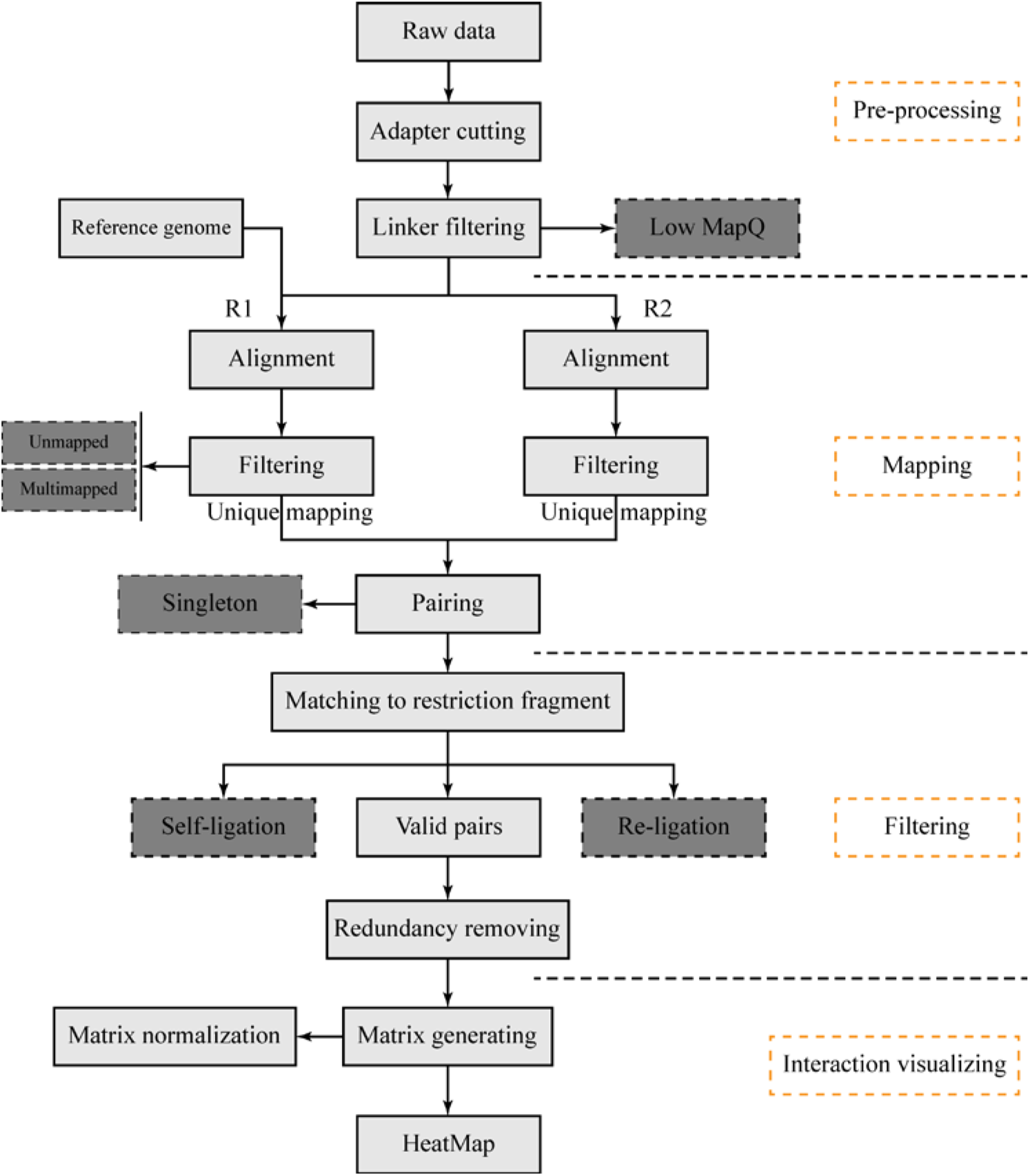
The schematic pipeline for DLO Hi-C Tool. Four separate modules of DLO Hi-C Tool to process DLO Hi-C data.

### Data pre-processing

#### Adapter cutting

The effective sequence from the DLO Hi-C construct is about 80-bp. With the present sequencing capacity, single-end sequencing can get the full-length of the constructs. If paired-end sequencing protocol is applied, only the first sequenced reads, which contain the whole effective sequences, are kept for further analysis. In the current sequencing protocol, indexed adapters are generally added in the end of the reads (Fig. 2). At the beginning of data processing, the adapters need to be clipped from the raw reads (Fig. 1). The input file can be FASTQ or gzip’d FASTQ format. In order to get accurate sequences of interactions, the accuracy of the start position of adapter is very critical. At this step, mafft [11] is used to recognize the sequences of adapter. Multiple sequence alignment is conducted on 100 reads, and the adapter sequence is recognized based on the alignment results. The program will count the base frequency in each position from alignment result and then estimate the most probable base in each position. Then, the adapter sequences were detected and cut from the raw reads. If the user don’t want cut adapter or want to use given adapter sequence, they can set the parameter “AdapterSeq” blank or the sequence want to used.

**Fig. 2.**
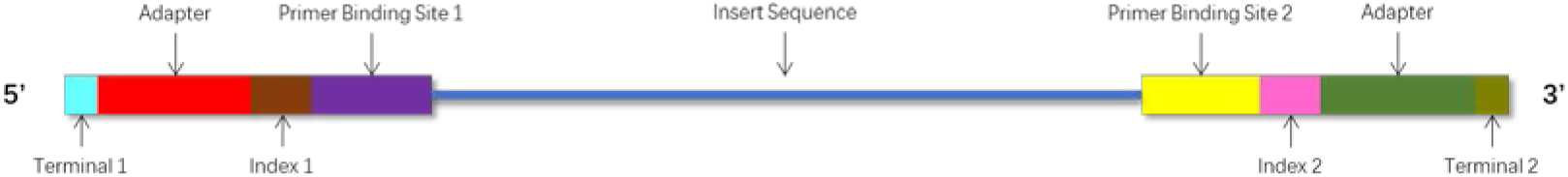
The composition of Illumina paired end sequencing reads.

#### Linker detection and filtering

Because of the specific linker sequences, it is easy to distinguish the interaction fragments from genomic DNA. Besides, the linkers were also designed to estimate the ratio of random ligation noise. The convenience to estimate the ratio of random ligation noise with linkers in DLO Hi-C library makes a contribution to its advantages. DLO Hi-C Tool will make a combination of all possible linkers according to the half-linker sequences.

There are 4 combinations (AA, BB, AB and BA) of half-linkers in DLO Hi-C and 1 combination (AA) of half-linkers in in situ DLO Hi-C library. The length of linkers is about 40-bp, which is located in the middle of the sequenced reads. The combinations of linkers (two half-linkers) are first aligned to the adapter clipped reads to get the precise position of linkers (Fig. 1). The result file of this step contains the information of every reads, such as the sequence of the left-linker, the start index of the linker, the end index of the linker, the sequence of the right-linker, the start position of adapter, linker type, mapping quality, read label, read sequence, sequence orientation and sequencing quality. Then the reads with linkers will be separated into 4 different files (DLO Hi-C) or 1 file (in situ DLO Hi-C) based on linker types for the following steps.

When lacking the information of linker sequence, it is essential to detect the linker sequence automatically. With this consideration, a module is designed to recognize the linker sequence and it is integrated into the main process of DLO Hi-C Tool. This module can detect not only the linker sequence, but also the restriction enzyme cutting site used in the experiment. As far as we know, it is the first pipeline that can detect linker and enzyme cutting site automatically. Without the information of linker of enzyme, users can set the ’HalfLinker’ or ’Restriction’ value to an empty string in the pipeline.

### Mapping

#### Mapping the reads to the reference genome

The interaction products can be obtained according to the start and end indexes of linkers. Read pairs are aligned to the corresponding reference genome separately by Burrows-Wheeler Alignment (BWA) [12, 13] with aln algorithm. The SAI file is produced with 0 maximum edit distance and other default arguments. Not only is it time consuming to convert SAI file to SAM file, but also the conversion cannot be processed in a parallel mode. In order to reduce time spent in alignment, the FASTQ file is split into several chunks, and the alignment of each chunk is conducted on each individual thread. After aligning to the reference genome and converting (Fig. 3a), the mapping results are merged. Besides, if the sequence reads are long (more than 70-bp), the alignment of long sequence is supported by setting ’ReadsType’ as ’long’ for mem algorithm of BWA. Based on the mapping score, reads are divided into three categories, uniquely mapping reads with the mapping score more than 20, multiple locations with the score less than 20 and unmapped reads with the score of 0. If mem algorithm of BWA is applied for alignment, the mapping score cutoff is set to 30.

**Fig. 3.**
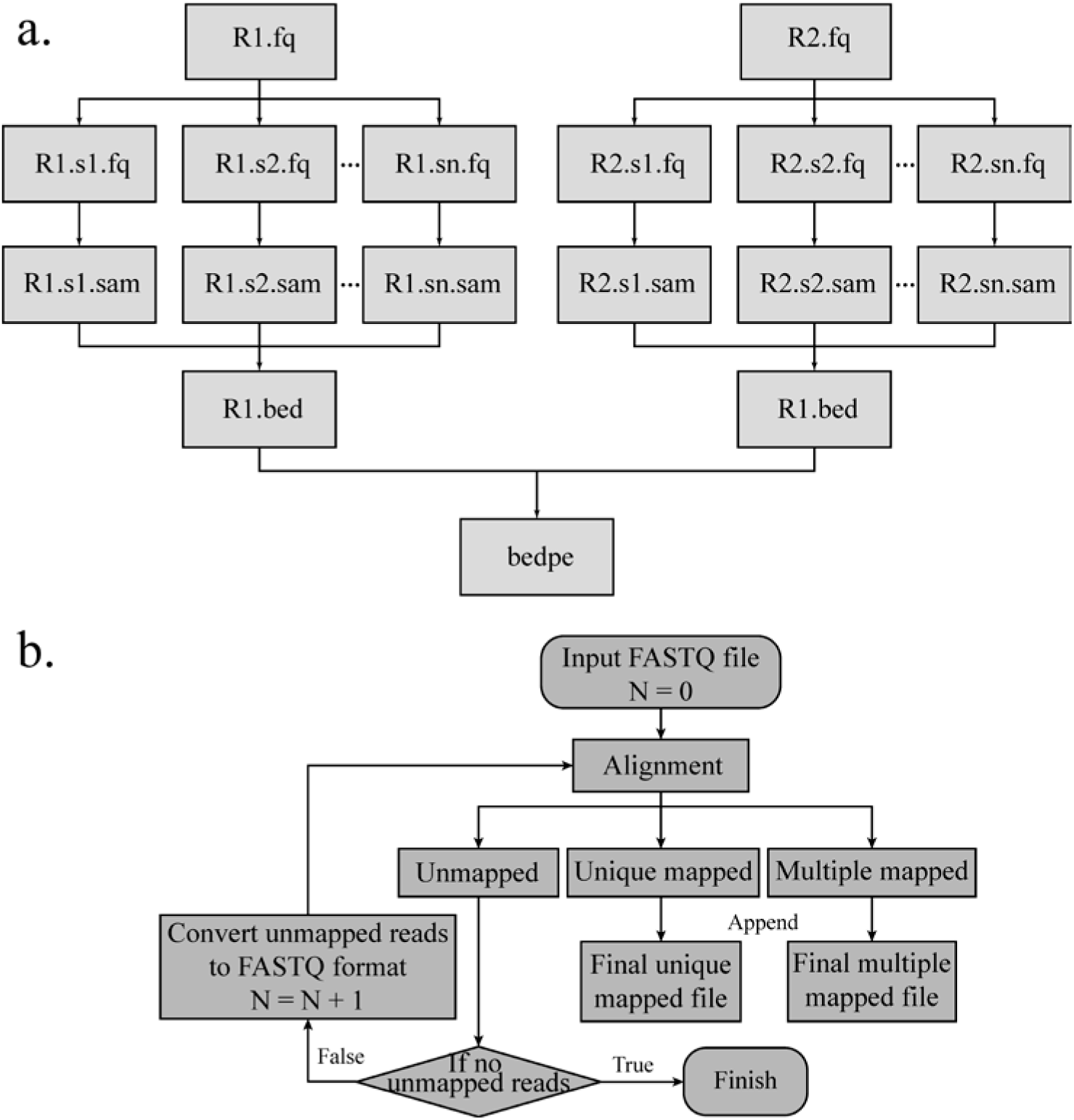
Flow chart of mapping and iterative mapping. a, Raw data are split into several chunks and the alignment results of different chunks will be merged after pairing. b, The flowchart shows the detail steps of iterative mapping, N represents the maximum edit distance of BWA.

#### Iterative mapping

DLO Hi-C Tool supports an additional mapping method named “Iterative mapping”. User can choose whether to use the method when mapping. There is a parameter “n” in BWA aln algorithm and the value represents the max mismatch base allowed. In the beginning (Step 1), the maximum edit distance N will be set to 0 (Fig. 3b). After mapping, all reads will be classified into 3 categories, mapped, unmapped and multi-mapped. In Step 2, the reads corresponding to unmapped category in the previous step will do alignment again with N = 1, and the alignment results will be classified into 3 categories. In Step 3, the reads in unmapped category from Step 2 will be mapped with one more mismatch allowed, which will be repeated with iterative mapping until no unmapped reads. In the end, all reads of “mapped” and “unmapped” categories will be retained and merged. Iterative mapping can utilize the reads which include SNP (single nucleotide polymorphism) or sequencing error, so that increasing the unique mapping ratio.

### Noise reduction and paired-end reads classification

#### Classification of paired-end reads based on alignment

Some of the paired-end reads are invalid interactions, such as re-ligation and self-ligation reads [10]. In addition, there are some duplicate reads produced by polymerase chain reaction (PCR). The combination of different linkers is used to estimate the ratio of random collision which can’t be obtained from the real biological complexes. If the two ends of reads correspond to two adjacent restriction fragments, the reads are called as re-ligation reads. These reads are ligated with the same linker after digestion. If two ends of a DNA fragment are ligated with one linker after digestion, the corresponding reads are self-ligation ends (Fig. 4).

**Fig. 4.**
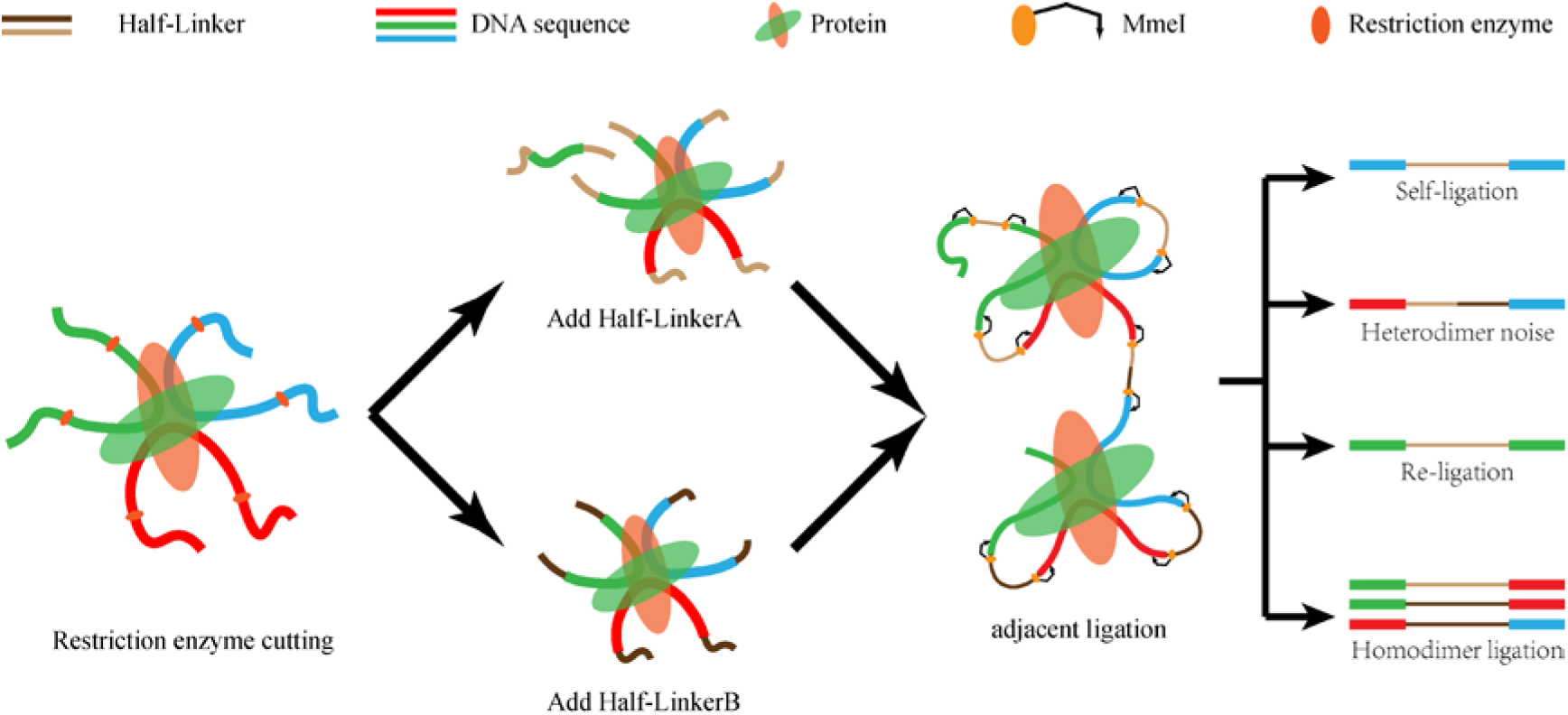
The sketch map of self-ligation and re-ligation.

#### Duplication removing

The duplicates with the same strands and positions of two ends produced by PCR need to be filtered out before interaction calling. Too many duplicates indicate high duplication noise. In order to reduce the effect of duplicate sequencing, we consider the reads with both ends mapped within 2 bp difference respectively are duplicate reads. Then we will remain only one read of these reads for further analysis.

#### Interaction matrix converting and normalization

The resolution is the bin size used in the contact maps. The length of restriction fragment is the extreme resolution of Hi-C library. The details differ for each resolution. We could get more information from high resolution (small bin size) map, but it needs more sequencing depth and is time-consuming in computation. The processing time is shorter for low resolution (big bin size), but it may loss some details. We should make a balance between the computing resources and map details. Users can define the values of the resolution in the configuration file to construct the interaction heatmap.

#### Statistics Reports

DLO Hi-C Tool will produce a Hyper Text Markup Language (HTML) report with the combination of the results, including the parameters used in the pipeline, the information of adapter and linker sequences and the running time of all steps. We also provide a Graphical User Interface for users to set the parameters conveniently.

#### Configuration file

There are two configuration files in DLO Hi-C Tool, commonly used parameters and advanced parameters. The commonly used parameters need to be modified for each DLO Hi-C library, and the advanced parameters (as optional parameters) can be kept as the default values. The format of configuration file contains three parts: the name of argument, equal sign, and the value of parameters. User can set the value of optional parameters blank if they want to use default value. Program will load advanced configure first then load commonly configure, so that if a parameter appear in two file, program will use the value in commonly configure file.

## Results

DLO Hi-C Tool was used to detect the linker of previously published DLO Hi-C data [10]. The test data includes 1 DLO Hi-C library of K562 cell line, 3 replicates of in situ DLO Hi-C library of K562 cell line and two replicates of in situ DLO Hi-C library of THP-1 cell line. The disk space of test files is listed in Table 1.

**Table 1.** The disk space of test file

| Cell line | Enzyme | Dataset | Raw reads | File size |
| --- | --- | --- | --- | --- |
| K562 | HindIII | K562-HindIII | 228,257,962 | 77G x1 |
| K562 | MseI | K562-MseI-1 | 191,931,665 | 64G x1 |
| K562 | MseI | K562-MseI-2 | 208,310,948 | 70Gx1 |
| K562 | MseI | K562-MseI-3 | 262,150,931 | 88G x1 |
| THP-1 | MseI | THP-1-MseI-1 | 402,394,454 | 145G x1 |
| THP-1 | MseI | THP-1-MseI-2 | 431,193,228 | 156G x1 |

The first column is the cell line used. The second column corresponds to the restriction enzyme used in the experiment. The third column is the labels of the data sets. The fourth column shows the number of raw reads in Read1 file. The fifth column is the disk space of raw data in Read1 file with FASTQ format.

### Pre-processing

Pre-processing includes 5 main steps: linker detection, adapter detection, linker alignment, separation and linker filtering. Table 2 shows the results of the linker and adapter detection by DLO Hi-C Tool. The types of restriction enzyme and linker sequence were consistent with the experiment. The read length of test dataset is paired-end 2 x 150 bp. Taking into account the quality of sequencing, we only kept the R1 end for subsequent analysis.

**Table 2.**
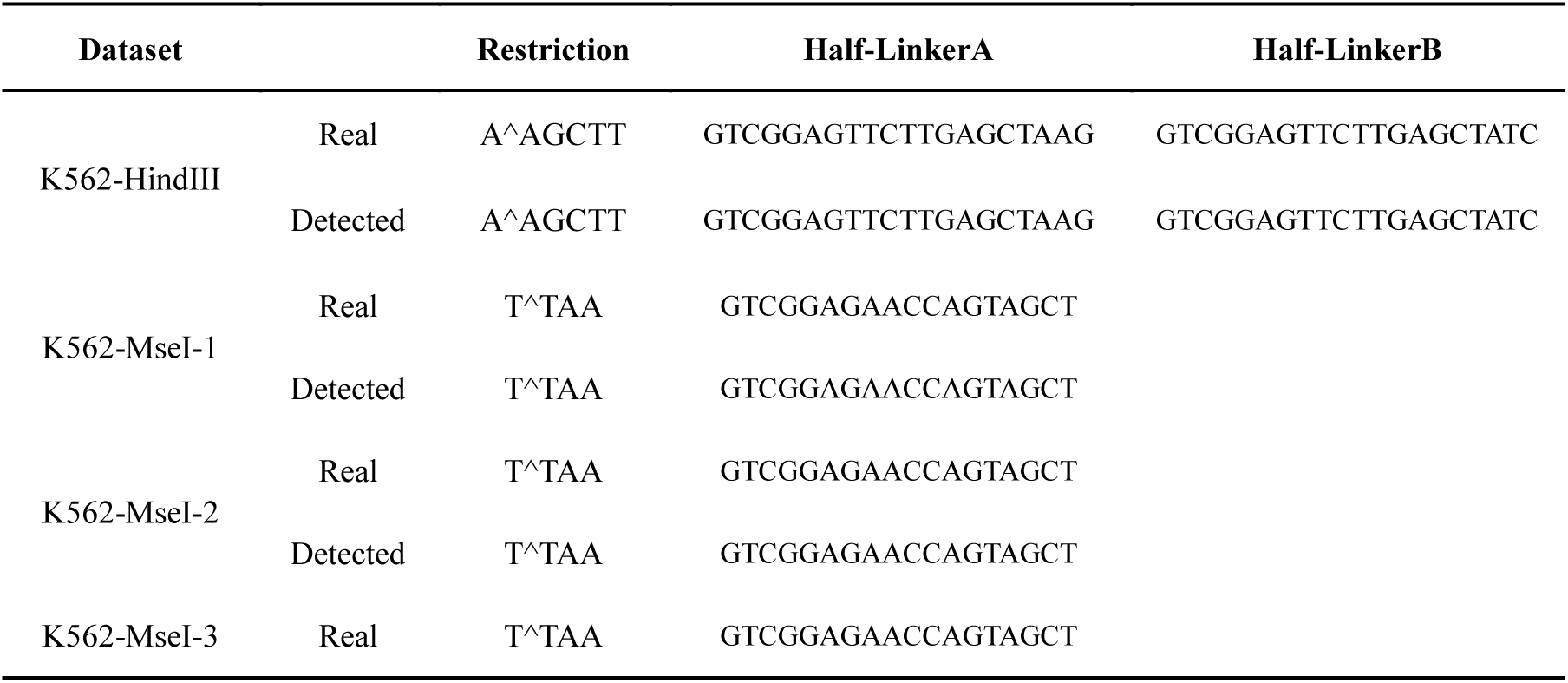
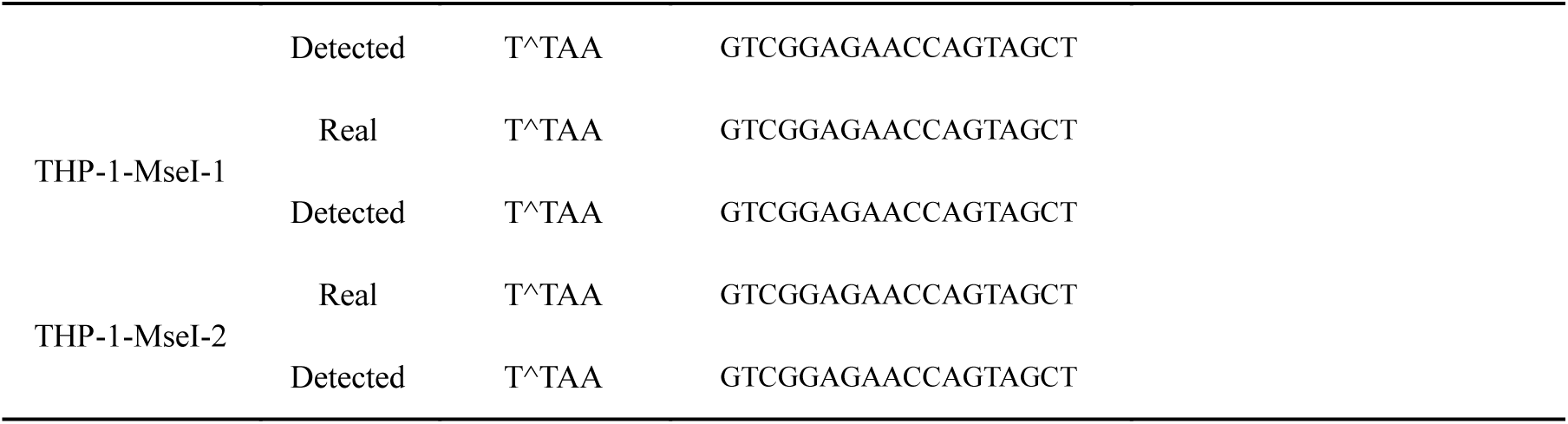
DLO Hi-C Tool performance on enzyme and linker detection

We refer to the algorithm of linker mapping in ChIA-PET Tool[14], and make some modifications to enable multi-threads execution. The parameter used in the alignment process is as follows, MatchScore: 1, MisMatchScore: −1 and InDelScore: −1. The linkers can be aligned to 10 million reads by DLO Hi-C Tool in 1 minutes and 42 seconds with 16 threads, which is much more efficiently than shell pipeline (Table 3). The results of pre-processing were saved in 01.Preprocess directory. It includes 2 FASTQ files and 1 txt file.

**Table 3.**
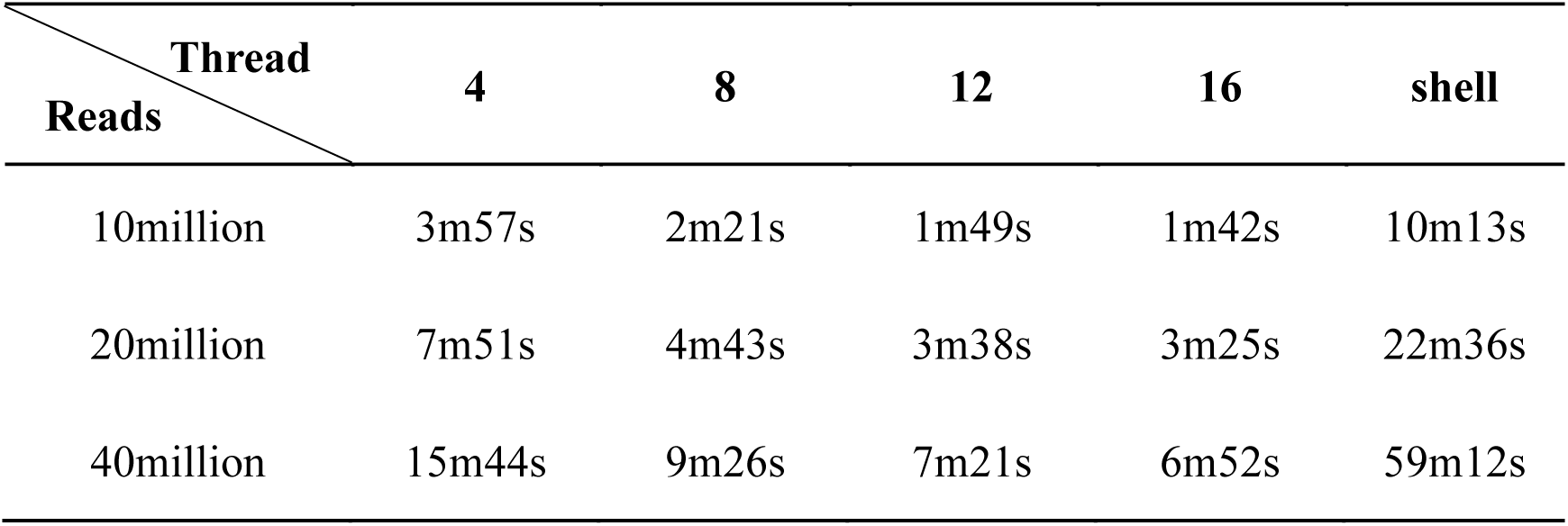
Running time of linker filtering (K562-MseI-1)

| Thread<br>Reads | 4 | 8 | 12 | 16 | shell |
| --- | --- | --- | --- | --- | --- |
| 10million | 3m57s | 2m21s | 1m49s | 1m42s | 10m13s |
| 20million | 7m51s | 4m43s | 3m38s | 3m25s | 22m36s |
| 40million | 15m44s | 9m26s | 7m21s | 6m52s | 59m12s |

We have calculated the rate of the proportion of the sequence with linkers. Of all raw reads, the ratio of linker sequence ranges from 78.2%-88.8% (Fig. 5a).

**Fig. 5.**
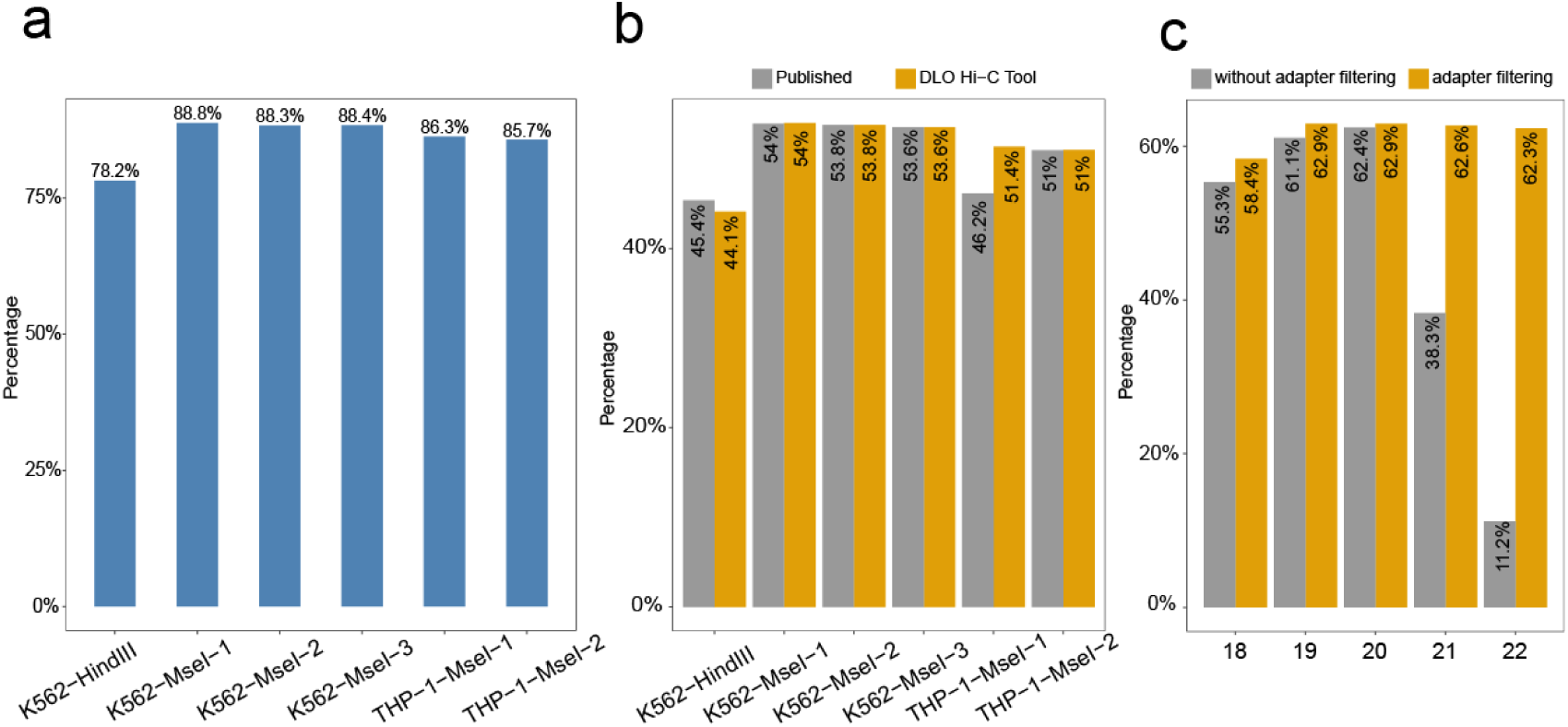
The ratio of linker sequence and uniquely mapping rate. a, The proportion of reads containing the linker to the raw reads. b, The uniquely mapping rate obtained from DLO Hi-C Tool without iterative alignment and published results of different samples. c, After adapter filtering or without adapter filtering, the change trend uniquely mapping rate as maximum read length increases.

### Alignment

BWA was used to align the linker filtered reads to the reference genome, and the default parameter is -t 1, -n 0. 20 was set as a cutoff score of unique mapping. The unique mapping ratio is similar to that of reported (Fig. 5b). The ratio ranges from 44.1%-54.0%.

Besides, we have tested the effect of adapter filtering on unique mapping by setting different maximum reads length (Fig. 5c). By comparing the percentage of unique mapping with or without adapter at different Maximum interception length, we found that the percentage of unique mapping are nearly the same when the interception length is shorter than 20 bp, but the percentage of unique mapping without adapter filtering is declined rapidly with the interception length increasing. Overall, the percentage of uniquely mapping with adapter filtering is relatively stable.

DLO Hi-C Tool was used to process DLO Hi-C and in situ DLO Hi-C datasets published by Lin et al [10]. We compared the computation time and the number of reads after every step. The numbers of reads are nearly the same between two pipelines of every step. The uniquely mapping rate and data utilization rate have a slight improvement on DLO Hi-C tool. With interactive alignment, the uniquely mapping rate had improved by 5% and the rate of data utilization had improved by 3% (Table 4).

**Table 4.** Comparative of different strategies.

|  | Lin et al[10] | Without iteration | With iteration |
| --- | --- | --- | --- |
| Data set | K562-MseI-1 | K562-MseI-1 | K562-MseI-1 |
| Digestion enzyme | MseI | MseI | MseI |
| Raw reads | 191,931,665 | 191,931,665 | 191,931,665 |
| Linker reads | 169,854,725 | 169,885,029 | 169,885,029 |
| Uniquely mapped reads | 103,549,004 | 103,642,564 | 111,025,113 |
| Non-redundant mapped reads | 96,795,788 | 95,573,098 | 102,242,616 |
| Inter-chromosomal contacts | 17,637,527 | 16,383,731 | 17,986,194 |
| Intra-chromosomal contacts | 79,158,261 | 79,189,367 | 84,256,422 |
| Intra-short-range contacts <= 5K | 18,740,658 | 18,663,657 | 19,826,034 |
| Intra-long-range contacts > 5K | 60,417,603 | 60,525,710 | 64,430,388 |

After alignment, DLO Hi-C Tool will give a statistics report. Fig. 6 gives an example of the alignment statistics part of the final report. Besides the version of BWA and the directory of genome file, the report also gives the detail information of the parameter and the result of alignment. It is convenient for users to compare the results with different parameters.

**Fig. 6.**
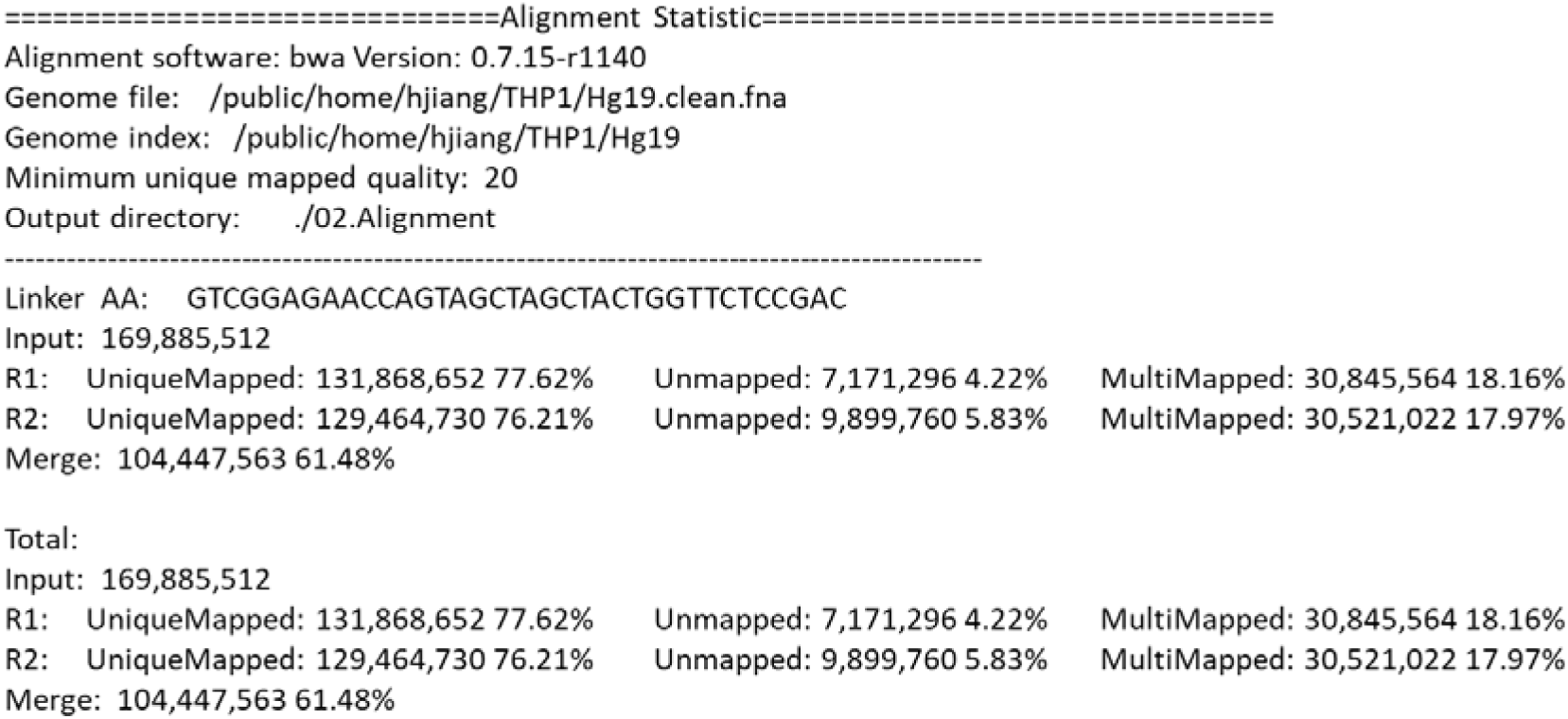
Statistics report of alignment.

### Duplicate Filtering

The statistics of noise redundancy is included in the final report (Fig. 7). For linker AA and linker BB sequence, it shows the detailed number and percentage of self-ligation, re-ligation, duplication, intra-chromosome, inter-chromosome, short range and long range reads separately. This report makes it easier for users to estimate the quality of DLO Hi-C library.

**Fig. 7.**
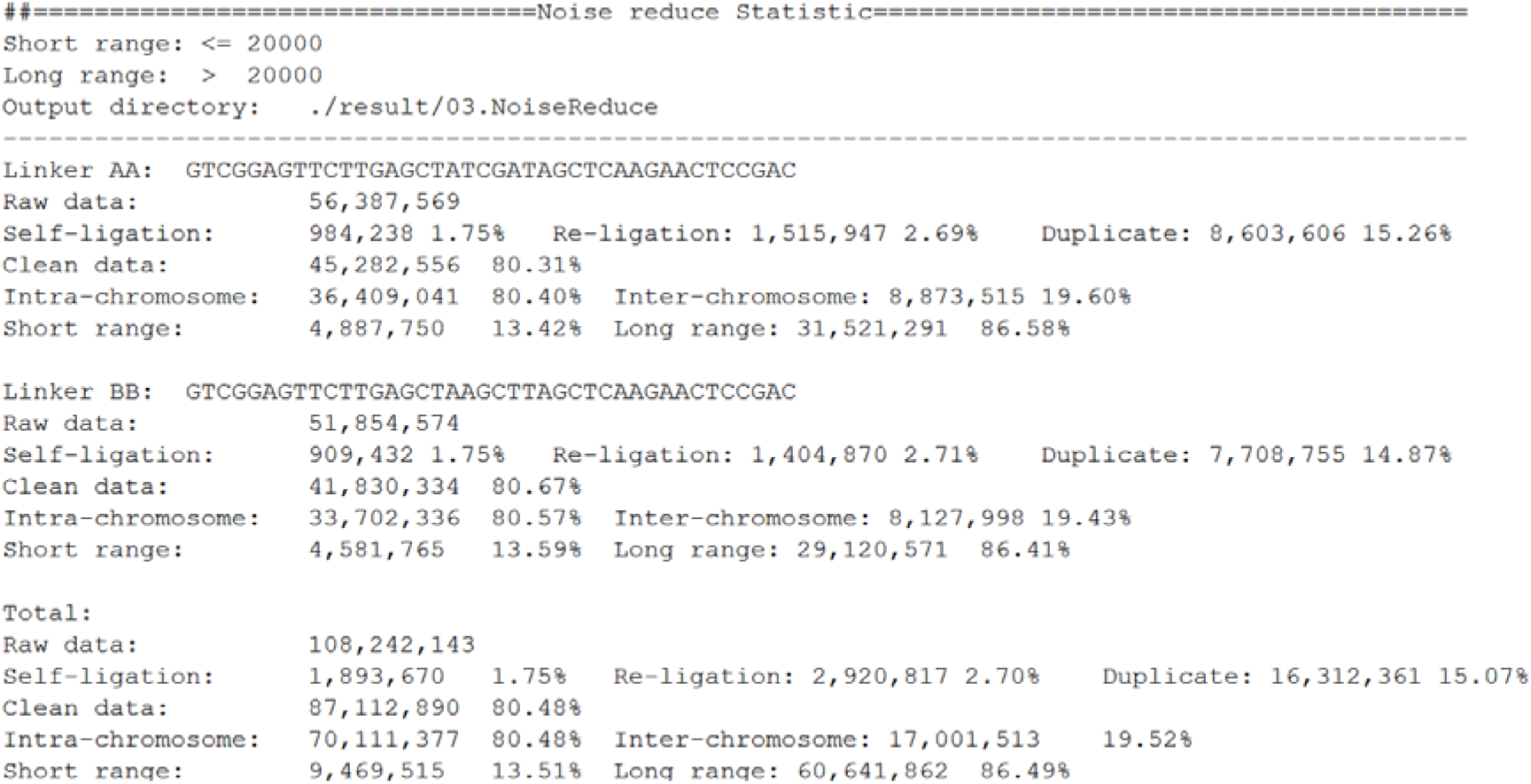
Statistics report of noise redundancy.

### Interaction Matrix

Then we compared the interaction heatmap of our results with published results. Fig. 8 shows the chromatin interaction heatmap obtained by the DLO Hi-C Tool and the published results. We can see a higher consistency of chromosome 4 from the heatmaps.

**Fig. 8.**
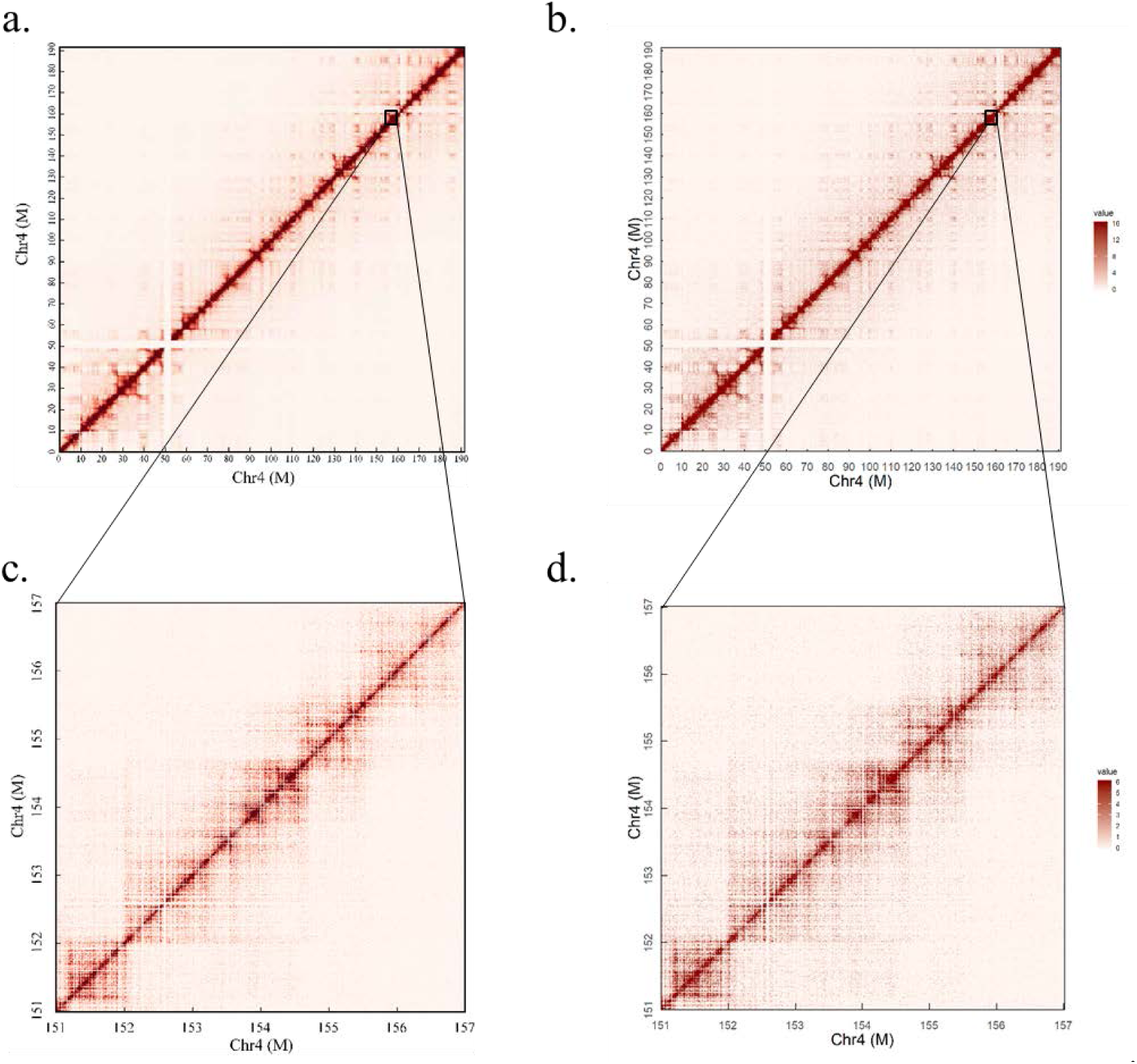
The interaction heatmap of chromosome 4 from K562 cell line. a and c, The published heatmap of the whole and local chromosome 4 of K562. b and d, The interaction heatmap of chromosome 4 and enlarged region generated by DLO Hi-C Tool.

With visual checking of the heatmaps, we found some potential chromosome translocations, as shown in Fig. 9. We confirmed the authenticity of these two translocations by the software hic_breakfinder and Hi-Ctrans [15, 16]. The heatmap generated by DLO Hi-C Tool will provide a visual theoretical basis for chromatin translocation calling.

**Fig. 9.**
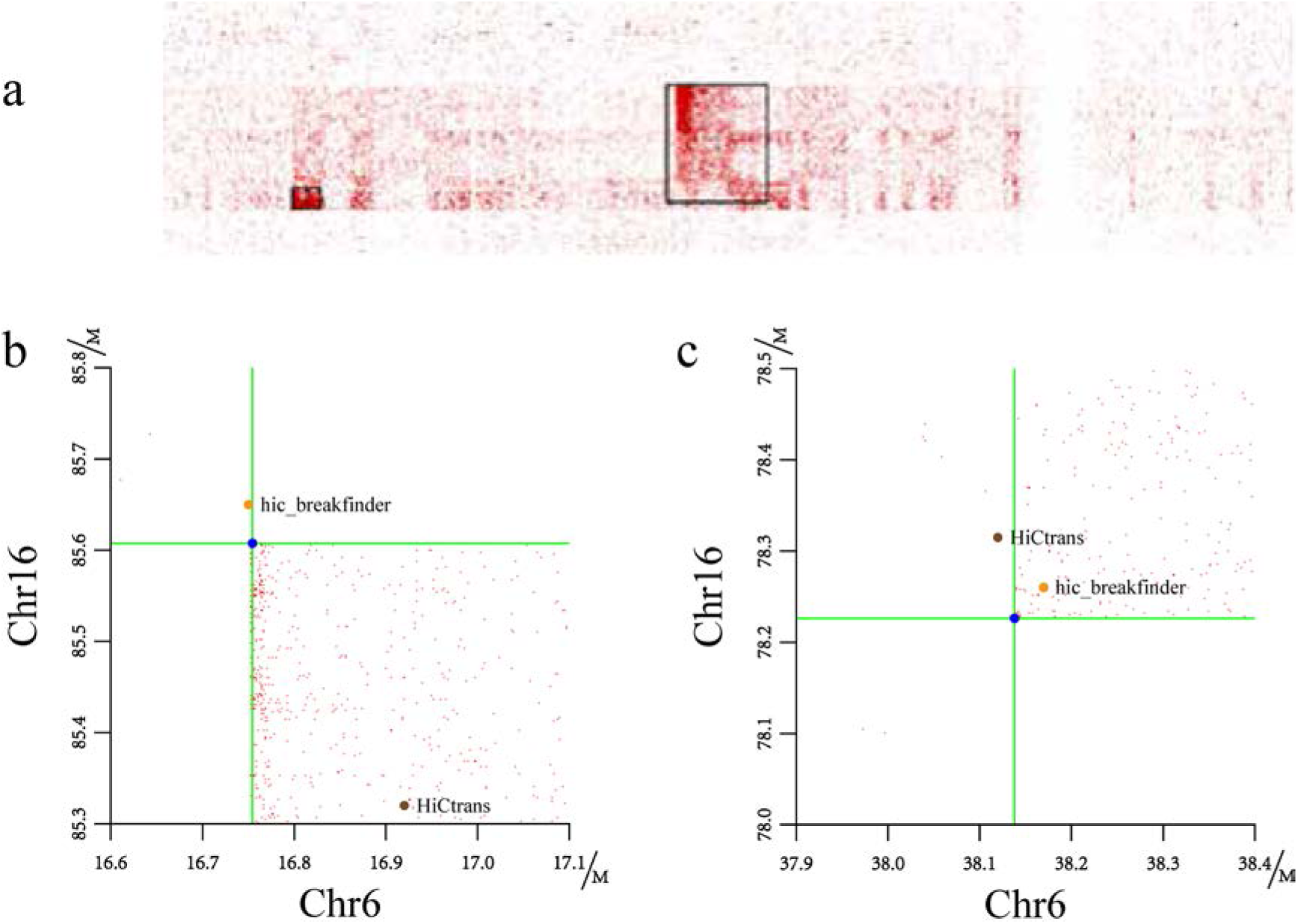
Translocation regions between chr6 and chr16. a, The translocation region obtained from heatmaps generated by DLO Hi-C Tool, and the translocations marked as black rectangles. b and c, the translocations find by hic_breakfinder and Hi-Ctrans.

### HTML report

In the final HTML format report, users will visually see the specific information of the entire DLO Hi-C library (Supplementary 1). The report mainly shows five aspects: Running information, Linker filtering, Alignment, Noise reducing and Matrix report.

The running information table indicates the detailed parameters used in the program. The linker filtering section contains basic statistics, base distribution in adapter detection, linker alignment score distribution and tag length distribution. From this part, it is easy to know the percentage and mapping details of linker sequences. The basic statistics of alignment gives the information of the reads number and percentages of different categories. In the basic statistics of the noise reducing section, we can get the number and proportion of the sequence of noise from different sources. Also the number and proportion of long and short range interactions have been list in the table. Besides, the statistics of position and negative chain and the interaction distribution are concluded in the noise reducing part. The matrix report part displays the interaction heatmap of whole genome and the running time of each step of the entire program.

## Conclusion and discussion

In this paper, we developed DLO Hi-C Tool for DLO Hi-C data analysis. By processing the data from the article of DLO Hi-C experiment, we demonstrated that DLO Hi-C Tool can get the results similar to those from the original shell scripts and it is more efficient. We have compared the features of DLO Hi-C Tool with the original shell scripts and found that DLO Hi-C Tool have more advantage than Shell script (Table 5). DLO Hi-C Tool supports compressed files as input, and have less software independence. DLO Hi-C needs lesser input information so that it is easier to use than Shell script. DLO Hi-C Tool can process data from any species without modifying any code. DLO Hi-C Tool will consume much memory when many threads are used. So, the threads used need to set at a small number, if memory isn’t enough.

**Table 5.** Comparison of DLO Hi-C Tool with the original shell script

| Item | Shell script | DLO Hi-C Tool |
| --- | --- | --- |
| Input file format | FASTQ | Original or gz compressed FASTQ |
| Thread | Single | Multiple |
| Software dependence | BWA, SAMtools, perl, BEDTools, Python | BWA, mafft, python |
| Robustness | Bad | Good |
| Resume from break point | Unsupported | Supported |
| HTML report | No | Yes |
| Linker sequence | Required | Optional |
| Restriction sites | Required | Optional |
| Statistics information | Little | Comprehensive |
| Running time<br>(test data 10million) | 131 minutes | 23 minutes<br>(8 threads) |

## Supporting information

web report

python version

## Availability and requirements

Project name: DLO Hi-C Tool.

Project home page: https://github.com/GuoliangLi-HZAU/DLO-Hi-C-Tool (Java version). https://github.com/GangCaoLab/DLO-HiC-Tools (Python version).

Operating system(s): Linux.

Programming language: Java.

Other requirements: JRE 1.8 or higher; mafft; BWA; Python 2.7; Python package: Matplotlib, SciPy, argparse and NumPy.

License: GPL license.

Any restrictions to use by non-academics: license needed.

### Abbreviations

3C: **C**hromosome **C**onformation **C**apture
3D: Three-Dimensional
BWA: Burrows-Wheeler Alignment
ChIA-PET: Chromatin Interaction Analysis by Paired-End Tag Sequencing
DLO Hi-C: Digestion-Ligation-Only Hi-C
HTML: HyperText Markup Language
PAGE: PolyacrylAmide Gel Electrophoresis
PCR: Polymerase Chain Reaction
SNP: Single Nucleotide Polymorphism
TADs: Topologically Associating Domains

## Declarations

### Ethics approval and consent to participate

Not applicable.

### Consent for publication

No applicable

### Availability of data and materials

The data used in this project were from NCBI GEO with accession number GSE89663.

### Competing interests

The authors declare that they have no competing interests.

### Funding

This research was funded by Natural Science Foundation of China (grant number 31771402 to G.L) and the Fundamental Research Funds for the Central Universities (grant number 2662018PY025 to G.C. and 2662017PY116 to G.L.).

### Authors’ Contributions

Study conceived by: P.H., D.L., G.C. and G.L.. Wrote the paper: P.H., H.J., W.X. and G.L.. Revised the paper: all authors. Pipeline testing: H.J., W.X. and Q.X..

## Acknowledgments

We are grateful for the support of our colleagues in testing DLO Hi-C Tool, particularly Muhammad Muzammal Adeel.

## Notes

#### Summary of Updates

author order change

## References

1. Gorkin DU, Leung D, Ren B: The 3D Genome in Transcriptional Regulation and Pluripotency. Cell Stem Cell 2014, 14(6):762–775.

2. Dekker J, Rippe K, Dekker M, Kleckner N: Capturing chromosome conformation. Science 2002, 295(5558):1306–1311.

3. Cullen KE, Kladde MP, Seyfred MA: Interaction between transcription regulatory regions of prolactin chromatin. Science 1993, 261(5118):203–206.

4. Liebermanaiden E, Van Berkum NL, Williams L, Imakaev M, Ragoczy T, Telling A, Amit I, Lajoie BR, Sabo PJ, Dorschner MO: Comprehensive mapping of long range interactions reveals folding principles of the human genome. Science 2009, 326(5950):289–293.

5. Fullwood MJ, Liu MH, Pan YF, Liu J, Xu H, Mohamed YB, Orlov YL, Velkov S, Ho A, Mei PH: An oestrogen-receptor-α-bound human chromatin interactome. Nature 2009, 462(7269):58–64.

6. Ma W, Ay F, Lee C, Gulsoy G, Deng X, Cook S, Hesson J, Cavanaugh C, Ware CB, Krumm A: Fine-scale chromatin interaction maps reveal the cis-regulatory landscape of human lincRNA genes. Nature Methods 2015, 12(1):71–78.

7. Rao SS, Huntley MH, Durand NC, Stamenova EK, Bochkov ID, Robinson JT, Sanborn AL, Machol I, Omer AD, Lander ES et al: A 3D map of the human genome at kilobase resolution reveals principles of chromatin looping. Cell 2014, 159(7):1665–1680.

8. Jager R, Migliorini G, Henrion M, Kandaswamy R, Speedy HE, Heindl A, Whiffin N, Carnicer MJ, Broome L, Dryden N: Capture Hi-C identifies the chromatin interactome of colorectal cancer risk loci. Nature Communications 2015, 6(1):6178–6178.

9. Dixon JR, Jung I, Selvaraj S, Shen Y, Antosiewiczbourget J, Lee AY, Ye Z, Kim A, Rajagopal N, Xie W: Chromatin architecture reorganization during stem cell differentiation. Nature 2015, 518(7539):331–336.

10. Lin D, Hong P, Zhang S, Xu W, Jamal M, Yan K, Lei Y, Li L, Ruan Y, Fu ZF et al: Digestion-ligation-only Hi-C is an efficient and cost-effective method for chromosome conformation capture. Nat Genet 2018, 50(5):754–763.

11. Katoh K, Misawa K, Kuma K, Miyata T: MAFFT: a novel method for rapid multiple sequence alignment based on fast Fourier transform. Nucleic Acids Res 2002, 30(14):3059–3066.

12. Li H, Durbin R: Fast and accurate long-read alignment with Burrows-Wheeler transform. Bioinformatics (Oxford, England) 2010, 26(5):589–595.

13. Li H, Durbin R: Fast and accurate short read alignment with Burrows-Wheeler transform. Bioinformatics (Oxford, England) 2009, 25(14):1754–1760.

14. Li G, Fullwood MJ, Xu H, Mulawadi FH, Velkov S, Vega V, Ariyaratne PN, Mohamed YB, Ooi HS, Tennakoon C: ChIA-PET tool for comprehensive chromatin interaction analysis with paired-end tag sequencing. Genome Biol 2010, 11(2):R22.

15. Dixon JR, Xu J, Dileep V, Zhan Y, Song F, Le VT, Yardımcı GG, Chakraborty A, Bann DV, Wang Y et al: Integrative detection and analysis of structural variation in cancer genomes. Nat Genet 2018, 50(10):1388–1398.

16. Chakraborty A, Ay F: Identification of copy number variations and translocations in cancer cells from Hi-C data. Bioinformatics 2018, 34(2):338–345.

