## Supplementary material for "DLO Hi-C Tool for Digestion-Ligation-Only Hi-C Chromosome Conformation Capture Data Analysis": web report

2019-8-21 21:23:27

### DLO-HiC


| | |
|---|---|
| Running information | 1 |
| Linker Filtering | 4 |
| Alignment | 1 |
| Noise reducing | 4 |
| Matrix report | 2 |


### Running information

#### Configure information

| | | | |
|---|---|---|---|
| Input file: | K562-HindIII-10M.fastq | Min reads length: | 33 |
| Output folder: | . | Max reads length: | 20 |
| Output prefix: | K562-HindIII-test | Match score: | 1 |
| Genome file: | /public/home/hjiang/THP1/Hg19.clean.fna | MisMatch score: | -1 |
| Genome index prefix: | /public/home/hjiang/THP1/Hg19 | InDel score: | -1 |
| HalfLinkerA: | GTCGGAGTTCTTGAGCTAAG | Resolution: | 1000000 2000000 |
| HalfLinkerB: | GTCGGAGTTCTTGAGCTATC | Thread: | 16 |
| Restriction: | A^AGCTT | | |

### Linker Filter

#### Basic statistics

| | | |
|---|---|---|
| Adapter Sequence: | Auto | |
| Total reads: | 10,000,000 | 100.00% |
| AA: | 3,679,890 | 36.80% |
| AB: | 128,838 | 1.29% |
| BA: | 130,804 | 1.31% |
| BB: | 3,915,612 | 39.16% |
| Ambiguous: | 2,141,467 | 21.41% |
| Output folder: | ./01.PreProcess | |

##### Base distribution in adapter detection

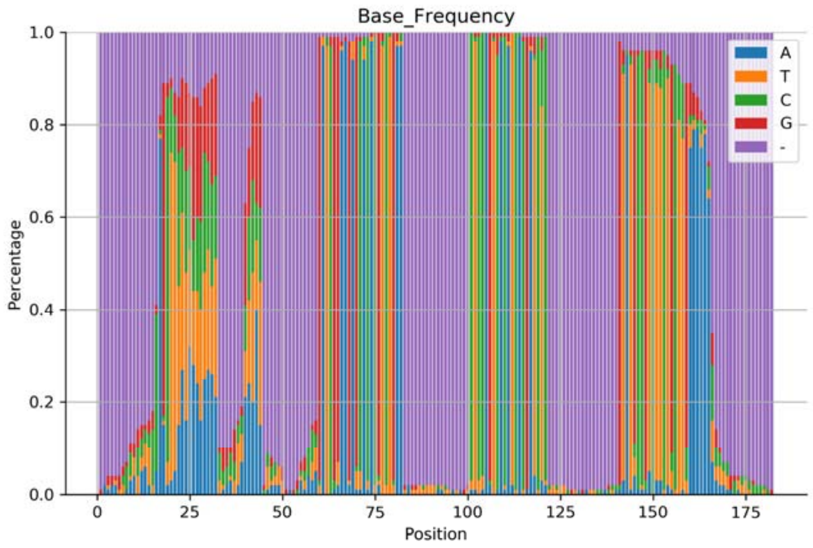


##### Linker alignment score distribution

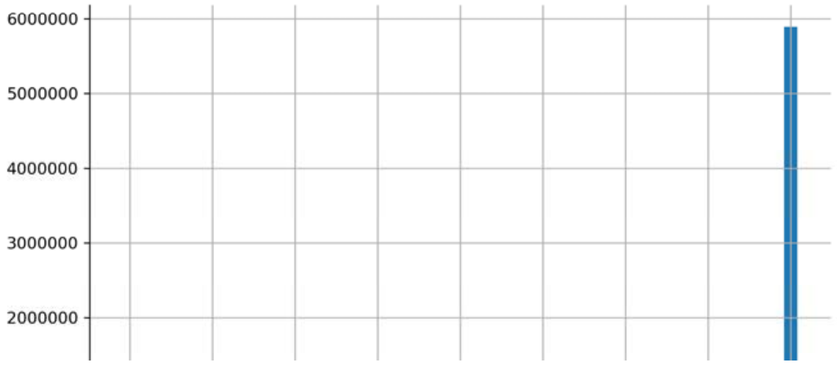

### DLO-HiC


Running information 1

Linker Filtering 4

Alignment 1

Noise reducing 4

Matrix report 2


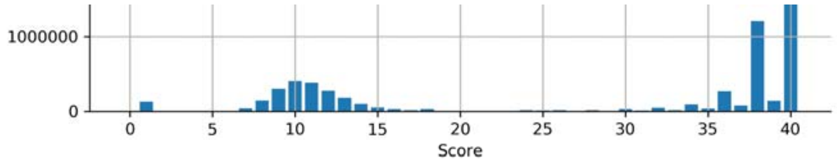


#### Tag length distribution

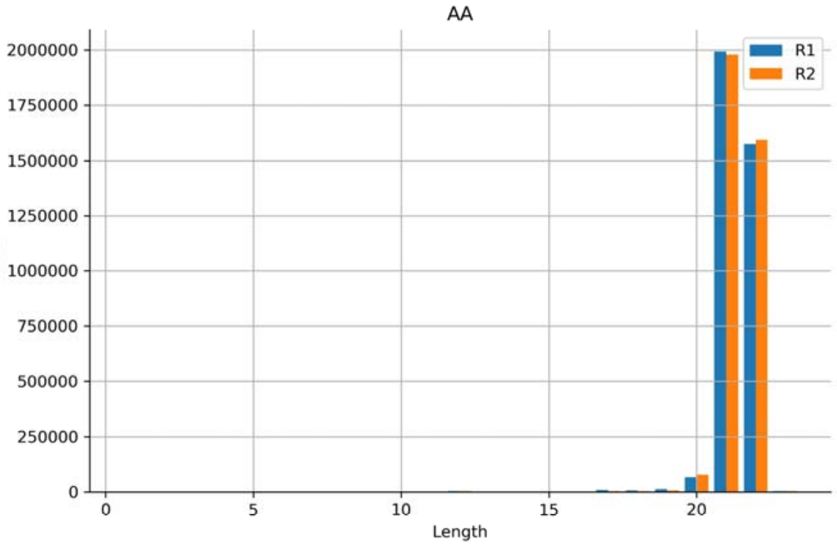


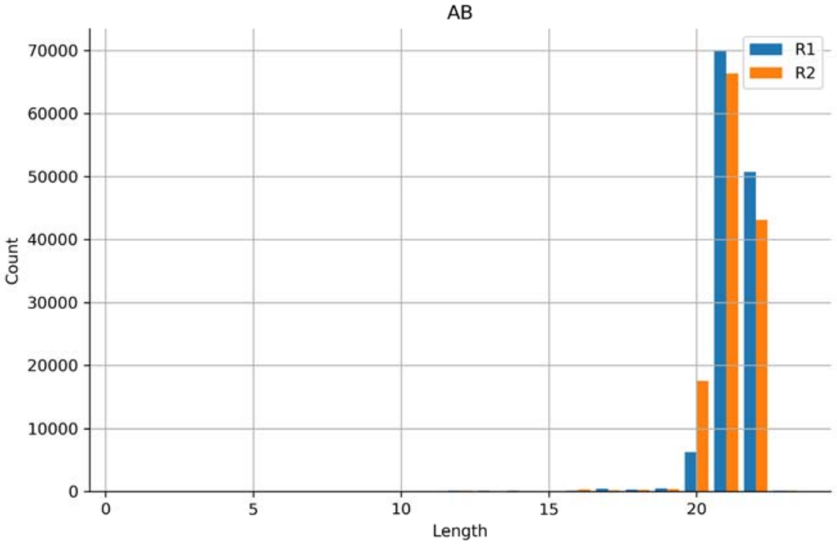


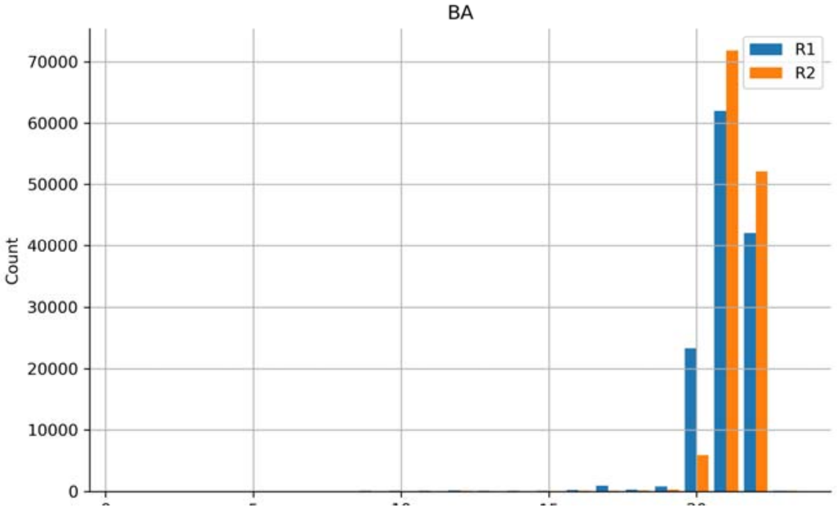

Running information 1
Linker Filtering 4
Alignment 1
Noise reducing 4
Matrix report 2


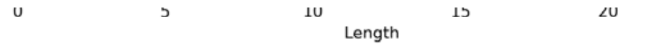


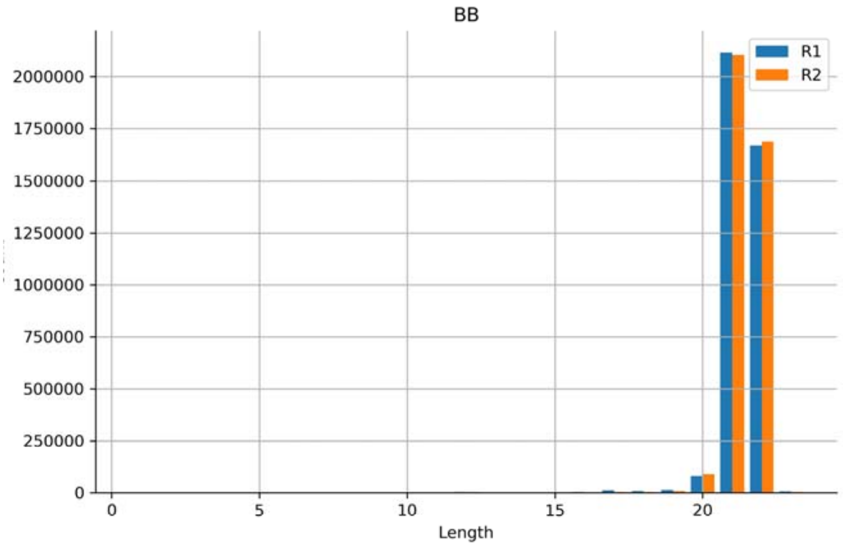


### Alignment

#### Basic statistics

##### Linker AA: GTCGGAGTTCTTGAGCTAAGCTTAGCTCAAGAACTCCGAC

| Item | Number | Percentage | Item | Number | Percentage |
|---|---|---|---|---|---|
| Fastq file R1: | 3,679,890 | 100.00% | Fastq file R2: | 3,679,890 | 100.00% |
| Unique map R1: | 2,977,042 | 80.90% | Unique map R2: | 2,966,807 | 80.62% |
| Multi map R1: | 702,848 | 19.10% | Multi map R2: | 713,083 | 19.38% |
| Unmap R1: | 0 | 0.00% | Unmap R2: | 0 | 0.00% |
| Merge: | 2,484,876 | 67.53% | | | |
| Output folder: | ./02.Alignment | | | | |

##### Linker BB: GTCGGAGTTCTTGAGCTATCGATAGCTCAAGAACTCCGAC

| Item | Number | Percentage | Item | Number | Percentage |
|---|---|---|---|---|---|
| Fastq file R1: | 3,915,612 | 100.00% | Fastq file R2: | 3,915,612 | 100.00% |
| Unique map R1: | 3,158,365 | 80.66% | Unique map R2: | 3,146,780 | 80.36% |
| Multi map R1: | 757,247 | 19.34% | Multi map R2: | 768,832 | 19.64% |
| Unmap R1: | 0 | 0.00% | Unmap R2: | 0 | 0.00% |
| Merge: | 2,628,236 | 67.12% | | | |
| Output folder: | ./02.Alignment | | | | |

### Noise reduce

#### Basic statistics

Input: 5,113,112

| Item | Number | Percentage | Item | Number | Percentage |
|---|---|---|---|---|---|
| Self-Ligation: | 90,137 | 1.76% | ReLigation: | 137,676 | 2.69% |
| Duplicate: | 429,963 | 8.41% | | | |
| Clean data: | 4,455,336 | 87.14% | | | |
| Intra-chrom: | 3,603,260 | 80.88% | Inter-chrom: | 852,076 | 19.12% |
| Short range: | 565,854 | 15.70% | Long range: | 3,037,406 | 84.30% |

#### Orientation-Position statistics

'+' and '-' represent the orientation of alignment, 's' means reads located in the 5' end of restriction fragment and 't' means reads located in the 3' end of restriction fragment

| | s,s | s,t | t,s | t,t |
|---|---|---|---|---|
| +,+ | 3,224 | 540,838 | 553,812 | 2,953 |
| +,- | 561,370 | 2,625 | 3,207 | 559,919 |
| -,+ | 560,128 | 3,258 | 2,801 | 561,004 |
| -,- | 3,743 | 541,837 | 552,135 | 2,482 |

DLO-HiC


Running information 1

Linker Filtering 4

Alignment 1

Noise reducing 4

Matrix report 2


#### Annotation statistics

#### Interaction distance distribution

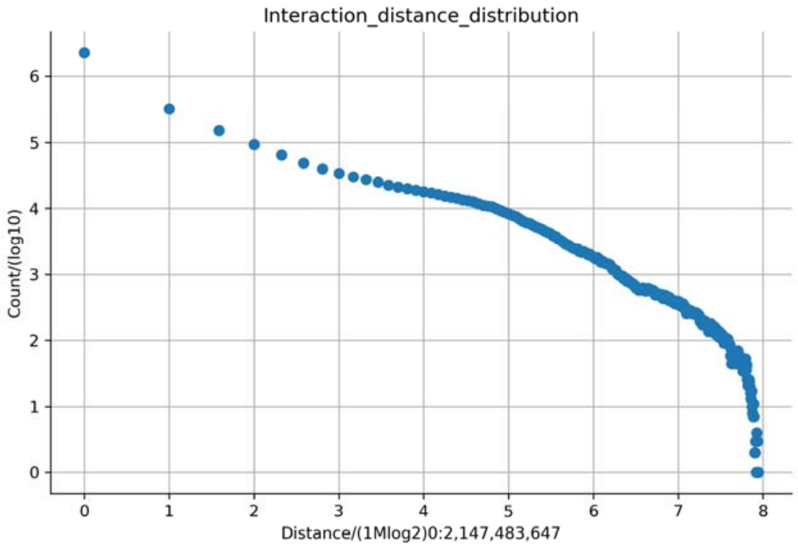


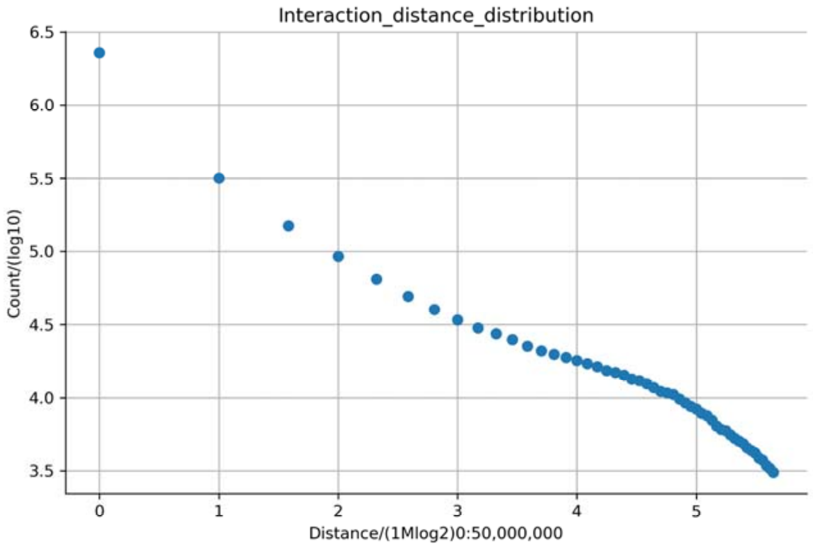


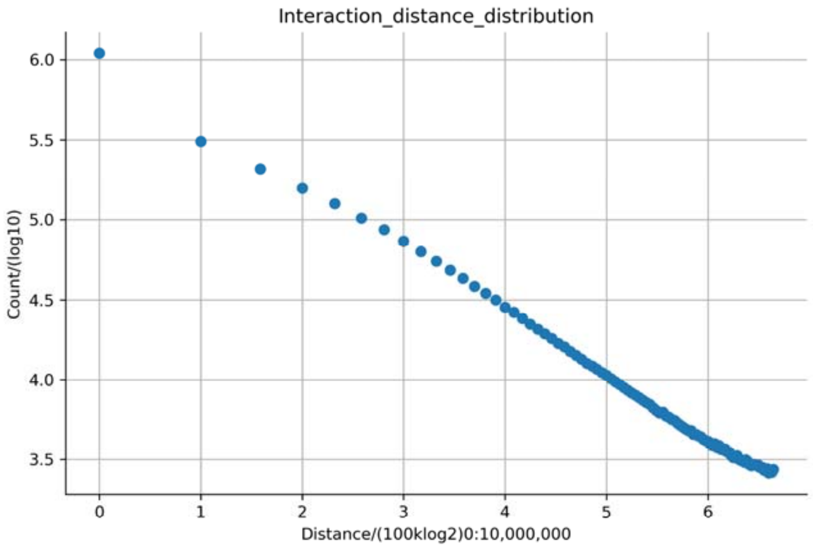


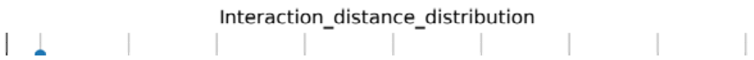

### DLO-HiC

Running information 1
Linker Filtering 4
Alignment 1
Noise reducing 4
Matrix report 2

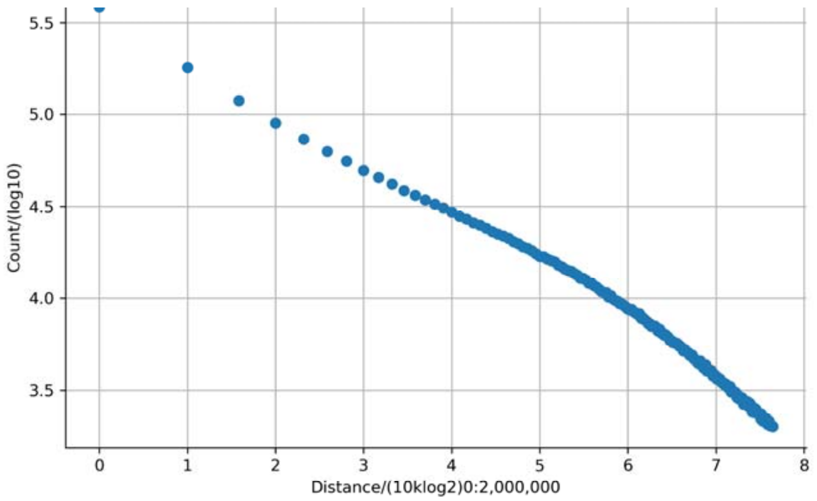


#### Matrix report

##### Basic statistics

###### Interaction heatmap

**Resolution 1,000,000**

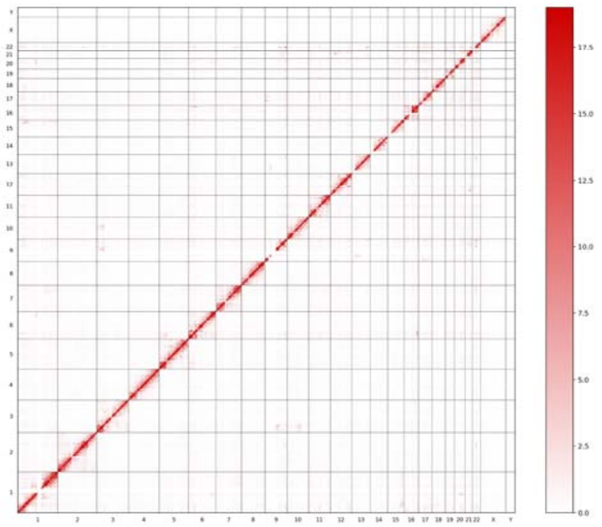

##### Running time

| Item | Value(h/m/s) |
|---|---|
| Start time: | Wed Aug 21 20:56:50 CST 2019 |
| Linker filtering: | 0H4M27S |
| Mapping: | 0H8M15S |
| Noise Reduce: | 0H1M5S |
| Create matrix: | 0H12M44S |
| Total: | 0H26M33S |

1. Please use Chrome, Firefox or Safari for better browsing experience.
2. Detailed explanation can be found at https://github.com/GuoliangLi-HZAU/DHat.
