## Supplementary material for "DLO Hi-C Tool for Digestion-Ligation-Only Hi-C Chromosome Conformation Capture Data Analysis": python version

Information about Python version

Implementation:
The strategies used in Python version DLO Hi-C Tool is basically same to the Java version. Tools used for each processing steps are wrapped into many command line tools, for user call them individually. And the pipeline is implemented with Snakemake[1], it's can solving the dependency relationships between each steps automaticlly, and compatible with compute clusters or batch systems like 'qsub'. So this pipeline can both run on single PC or a computer cluster for deal with huge amounts data. And the whole pipeline can be abort and resumed at any steps. The "Snakefile"(pipeline file) and the pipeline configuration template are generate using 'dlohic pipeline' command.
Because the Python interpreter is relative slow, for speed up the linker trim process, Cython [2]is used for performance optimization. And the Cython code used in semi-global pairwise alignment in linker and adapter match is come from the cutadapt[3]'s source code.

Software dependency and installation:
Python version DLO Hi-C Tool's is develop and tested depend on these software: Python(>=3.4), BWA(0.7.17), samtools(1.9), tabix(1.9), pairix(0.3.6), Cooler(0.8.3), mafft(v7.407)
User can install all dependency easily using conda package management system with provided instructions (https://github.com/GangCaoLab/DLO-HiC-Tools#requirements).
The Docker image is also provided (https://github.com/GangCaoLab/DLO-HiC-Tools#using-docker), by using the docker image user can using the DLO Hi-C Tool on different Operating Systems without any dependency and package installation.

Result file format :
The final interactions pairs are stored in 4DN-DCIC (Data Coordination and Integration Center)'s standard '.pairs' file (https://github.com/4dn-dcic/pairix/blob/master/pairs_format_specification.md). This file format can be indexed using the program pairix (https://github.com/4dn-dcic/pairix). Indexed file is random accessible, this feature allow fast fetch the interaction pairs within a specified genome region.
In this pipeline, the final interaction matrices will store to Cooler(4DN-DCIC standard)[4] or ".hic"(Aiden Lab)[5] format according to user's configuration. Both these file formats are support multiple-resolution matrices storage, matrix balancing and random access. Contact matrix in different resolution genome location can be quickly load into memory. It's convenient for the use of downstream analytic and visualization softwares based on these file formats, e.g. juicer tools[6], TADLib[7], juicebox[8], coolbox[9] etc.

Statistics report:
An example HTML report: https://nanguage.github.io/examples/DLO_HiC_Tools/test.html

Result:

Linker trim time:

| Thread<br>Reads | 4 | 8 | 12 | 16 | Shell Script |
|---|---|---|---|---|---|
| 10million | 2m27.971s | 1m42s | 1m42s | 1m42s | 10m13s |
| 20million | 4m44.673s | 3m24s | 3m14s | 3m23s | 22m36s |
| 40million | 8m24.267s | 6m22s | 6m23s | 6m35s | 59m12s |

Statistics of reads on MseI-1:

| | **Without iteration** | **With iteration** |
|---|---|---|
| Data set | K562-MseI-1 | K562-MseI-1 |
| Digestion enzyme | MseI | MseI |
| Raw reads | 191,931,665 | 191,931,665 |
| Linker reads | 169,538,630 | 169,538,630 |
| Uniquely mapped reads | 103,444,042 | 110,089,447 |
| Non-redundant mapped reads | 95,407,223 | 101,449,698 |
| Inter-chromosomal contacts | 16,351,769 | 17,862,785 |
| Intra-chromosomal contacts | 79,055,454 | 83,586,913 |
| Intra-short-range contacts <= 5K | 18,668,852 | 19,717,296 |
| Intra-long-range contacts > 5K | 60,386,602 | 63,869,617 |

Reference:


[1] Köster, Johannes, and Sven Rahmann. "Snakemake—a scalable bioinformatics workflow engine." *Bioinformatics* 28.19 (2012): 2520-2522.

[2] Behnel, Stefan, et al. "Cython: The best of both worlds." *Computing in Science & Engineering* 13.2 (2011): 31.

[3] Martin, Marcel. "Cutadapt removes adapter sequences from high-throughput sequencing reads." *EMBnet. Journal* 17.1 (2011): 10-12.

[4] Abdennur, Nezar, and Leonid Mirny. "Cooler: scalable storage for Hi-C data and other genomically-labeled arrays." *BioRxiv* (2019): 557660.

[5] Rao, Suhas SP, et al. "A 3D map of the human genome at kilobase resolution reveals principles of chromatin looping." Cell 159.7 (2014): 1665-1680.

[6] Durand, Neva C., et al. "Juicer provides a one-click system for analyzing loop-resolution Hi-C experiments." *Cell systems* 3.1 (2016): 95-98.

[7] Wang, Xiao-Tao, Wang Cui, and Cheng Peng. "HiTAD: detecting the structural and functional hierarchies of topologically associating domains from chromatin interactions." Nucleic acids research 45.19 (2017): e163-e163.

[8] Robinson, James T., et al. "Juicebox. js provides a cloud-based visualization system for Hi-C data." Cell systems 6.2 (2018): 256-258.

[9] Xu, Weize, et al. "CoolBox: a interactive genomic data explorer for Jupyter Notebook." BioRxiv (2019): 614222.
